# Antigen responsive CD4^+^ T cell clones contribute to the HIV-1 latent reservoir

**DOI:** 10.1101/2020.01.10.902155

**Authors:** Pilar Mendoza, Julia R. Jackson, Thiago Oliveira, Christian Gaebler, Victor Ramos, Marina Caskey, Mila Jankovic, Michel C. Nussenzweig, Lillian B. Cohn

**Affiliations:** Laboratory of Molecular Immunology, The Rockefeller University, New York, NY 10065; The Chan Zuckerberg Biohub, San Francisco, CA 94158; Howard Hughes Medical Institute, The Rockefeller University, New York, NY 10065; Department of Medicine, University of California, San Francisco. San Francisco, 94110

## Abstract

Antiretroviral therapy suppresses but does not cure HIV-1 infection due to the existence of a long-lived reservoir of latently infected cells. The reservoir has an estimated half-life of 44 months and is largely composed of clones of infected CD4^+^ T cells. The long half-life appears to result in part from expansion and contraction of infected CD4^+^ T cell clones. However, the mechanisms that govern this process are poorly understood. To determine whether the clones might result from, and be maintained by exposure to antigen, we measured responses of reservoir cells to a small subset of antigens from viruses that produce chronic or recurrent infections. Despite the limited panel of test antigens, clones of antigen responsive CD4^+^ T cells containing defective or intact latent proviruses were found in 7 out of 8 individuals studied. Thus, chronic or repeated exposure to antigen may contribute to the longevity of the HIV-1 reservoir by stimulating the clonal expansion of latently infected CD4^+^ T cells.

## Introduction

HIV-1 persists in memory CD4^+^ T cells including HIV-1, CMV, and influenza specific T cells (Douek et al., 2002; Jones et al., 2012; Demoustier et al., 2002; Hey-Nguyen et al., 2019). Following integration into the host genome, transcription usually leads to new virus production and cell death. However, HIV-1 can also become latent in a small number of infected CD4^+^ T cells, and these cells constitute a latent reservoir that is the principle barrier to HIV-1 cure.

A significant fraction of the circulating latent reservoir is composed of expanded clones of CD4^+^ T cells containing replication competent proviruses (50% or greater (Bui et al., 2017; Hosmane et al., 2017; Lorenzi et al., 2016; Simonetti et al., 2016; Reeves et al., 2018)). Although the origin of the clones and the mechanisms that govern their expansion are not known, longitudinal analysis indicates that they are dynamic, and change in size over time in individuals that are virologically suppressed on antiretroviral therapy (Wang et al., 2018).This dynamic may partially account for the longevity of the reservoir (Bruner and Cohn, 2019). Thus, understanding the basis for latently infected T cell clonal expansion is important for learning how to control and potentially eliminate the reservoir.

HIV-1 proviral DNA is enriched in HIV-1, CMV and influenza responsive T cells obtained from ART suppressed individuals, but whether or how this might be related to clonal expansion of T cells harboring latent viruses was not examined (Hey-Nguyen et al., 2019; Kristoff et al., 2019; Henrich et al., 2017; Douek et al., 2002; Demoustier et al., 2002; Jones et al., 2012) Here we report that expanded clones of CD4^+^ T cells containing replication competent viruses respond to antigenic stimulation with peptides derived from viruses that cause chronic infections.

## Results

To test the hypothesis that expanded clones harboring latent proviruses respond to foreign antigens, we exposed CD4^+^ T cells from antiretroviral therapy (ART) suppressed individuals (http://www.clinicaltrials.gov; NCT02825797; (Mendoza et al., 2018); NCT02588586; (Cohen et al., 2018); and NCT03571204, Supplemental Table 1) to overlapping peptide pools from either HIV-gag, CMV-pp65, or pooled peptides from CMV, EBV, influenza and tetanus toxin (CEFT). Staphylococcal enterotoxin B (SEB) and myelin oligodendrocyte glycoprotein (MOG), a self-protein, served as positive and negative controls respectively. Following overnight culture with HIV-gag, CMV-pp65, CEFT, or SEB, activated CD4^+^ T cells from 8 donors were purified by cell sorting based on expression of two or more activation induced markers (AIM+: CD69, PD-L1, and 4-1BB (Fig. 1a, b and Supplemental Fig. 1)). Total live CD4^+^ T cells were purified from parallel cultures stimulated with MOG to serve as unfractionated controls. As expected, there was little detectable response to the MOG self-antigen peptide pool, and all donors showed high level responses to SEB. In addition, responses to HIV-gag, CMV-pp65 and CEFT varied in magnitude among individuals (Fig. 1c and Supplemental Fig. 1).

**Figure 1.**
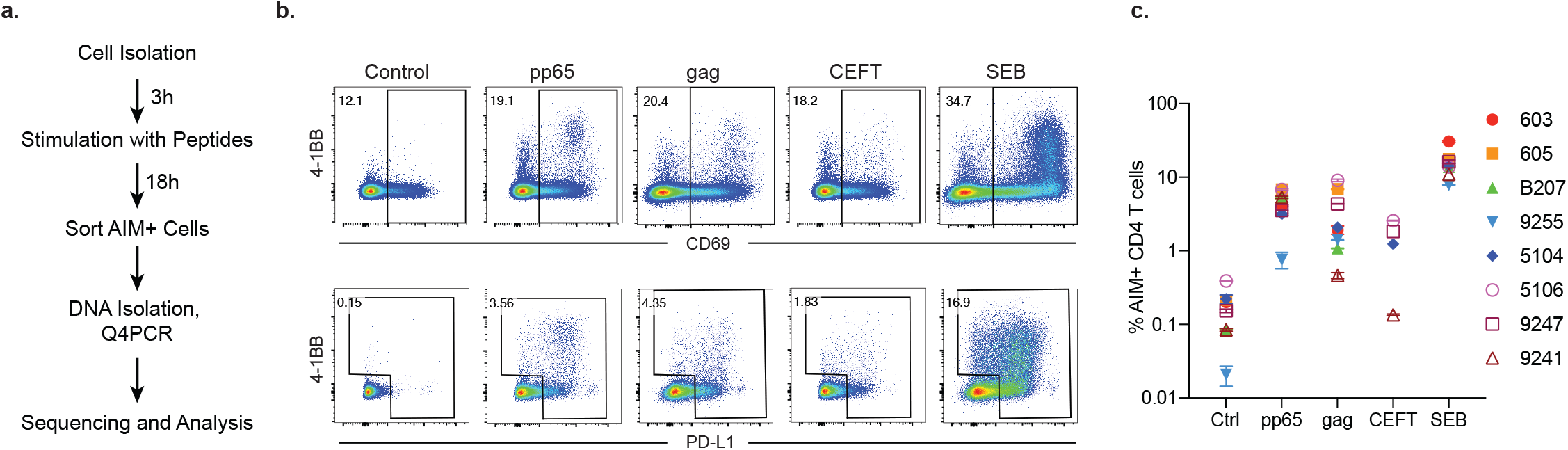
(a) Experimental overview. PBMCs were depleted of CD8^+^ T cells and rested for 3h before stimulation for 18h with peptide pools. Cells were then purified based on expression of activation induced markers (AIM) (CD69, PD-L1 and 4-1BB, Supplemental Figure 1). DNA was isolated from sorted cells, Q4PCR was performed, and sequenced viruses were assembled and analyzed. (b) Representative example of cell purification by AIM after stimulation with CMV-pp65, HIV-gag, CEFT (from 4/8 donors) or SEB by gating on CD69-positive cells followed by gating on PD-L1 positive or 4-1BB positive cells. Control CD4^+^ T cells cultured with MOG were purified on the basis of CD4 expression alone. Each AIM assay staining was performed twice on each donor. Numbers represent the percentage of total CD4^+^ T cells within each gate. (c) Frequency of AIM+ cells across all donors. Cells were processed and sorted as in (b).

To determine whether CD4^+^ T cells harboring HIV-1 proviruses are enriched among antigen or SEB responsive cells compared to the MOG control, integrated proviruses were enumerated and further characterized as intact or defective by combining quantitative PCR and next generation sequencing (Q4PCR, (Gaebler et al., 2019)) (Fig. 2a).

**Figure 2.**
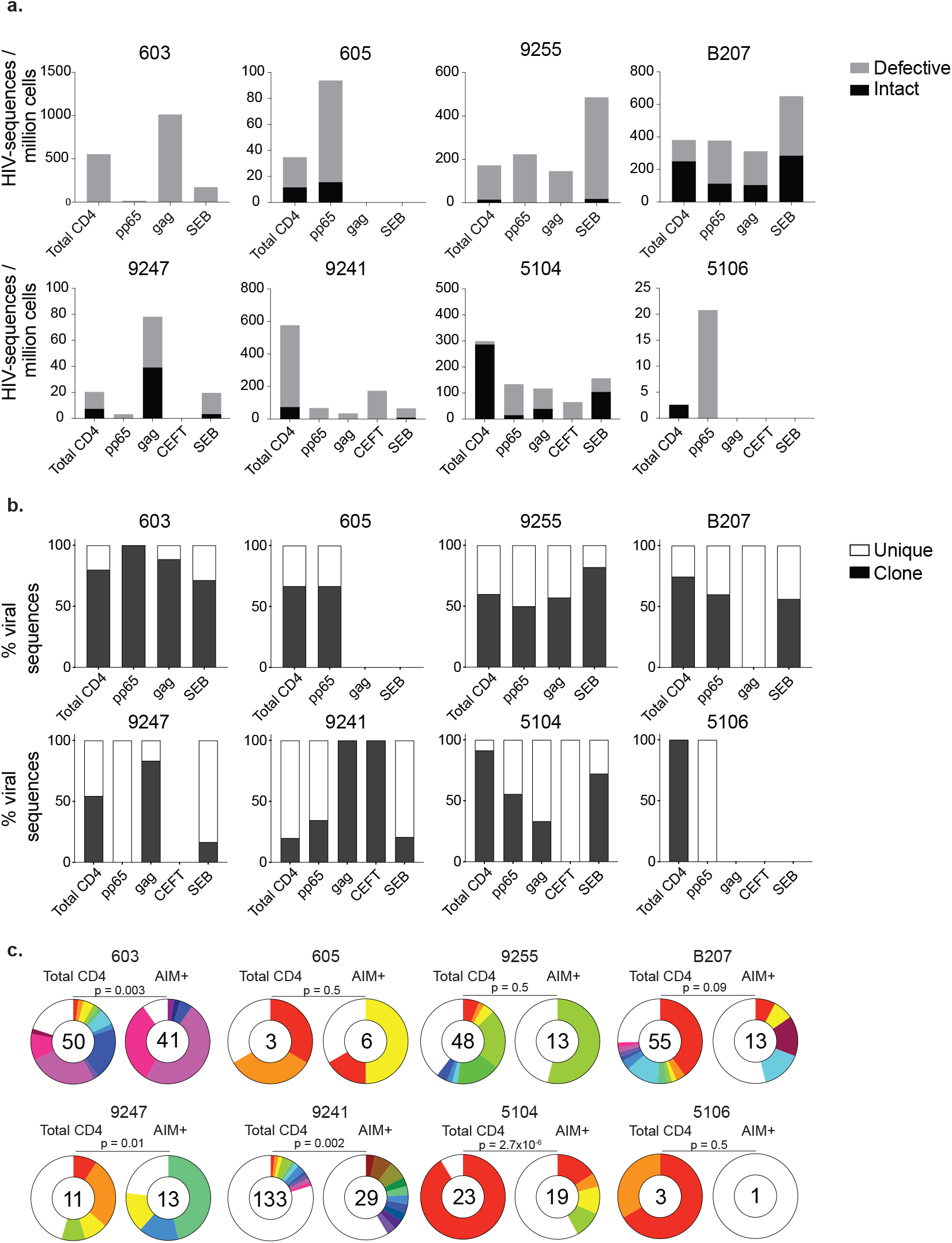
(a) Frequency of intact (black) and defective (gray) viruses characterized by Q4PCR per million AIM+ CD4^+^ T cells from 8 donors. (b) Frequency of virus sequences identified more than once (clonal) or only once (unique) among AIM+ cells from each donor. (c) Clonal distribution of Q4PCR-derived HIV-1 sequences from total CD4^+^ T cells (MOG control) and all antigen-reactive AIM+ CD4^+^ T cells combined, from 8 donors. Numbers in the inner circles indicate the total number of HIV-1 sequences analyzed. White represents sequences identified only once across all conditions (unique) and colored slices represent sequences identified more than once (clones). The size of each pie slice is proportional to the size of the clone. Asterisks denote a significant change in overall clonal distribution (* *P* ≤ 0.05; ** *P* ≤ 0.01; *** *P* ≤ 0.001, two-sided Fisher’s exact test).

The overall frequency of intact and defective proviruses contained within antigen responsive cells varied substantially among individuals. For example, proviruses were absent from CMV-pp65 specific CD4^+^ T cells in 603 and from HIV-gag specific CD4^+^ T cells in 605 or and 5106. In contrast, 1 in 1,000 HIV-gag responsive cells harbored a provirus in 603 (Fig. 2 and Supplemental Table 2). Individuals 603 and 9247 showed proviral enrichment in CD4^+^ T cells responding to gag of 2 and 4-fold, respectively, whereas in 605 and 5106 proviruses were 3 and 10-fold enriched in pp65 responsive CD4^+^ T cells (Fig. 2a). Finally, antigen responsive cells were relatively depleted of integrated proviruses in participants 9241 and 5104 (Fig. 2a). We conclude that HIV-1 proviruses can persist in CD4^+^ T cells that respond to CMV-pp65 and HIV-gag antigen with strong variation among infected individuals.

We analyzed all HIV-1 sequences across all groups to determine the fraction of viral sequences contributing to expanded clones of CD4^+^ T cells. Sequences found more than once were classified as derived from clones, while unique sequences were those identified only once. As expected (Bui et al., 2017; Cohn et al., 2015; 2018; Maldarelli et al., 2014; Lorenzi et al., 2016; Simonetti et al., 2016; Wang et al., 2018), clones of viral sequences were identified in all participants (Fig. 2b). The clonal distribution of HIV-1 sequences in antigen responsive (HIV-gag, CMV-pp65 and CEFT) was significantly different from the MOG control in 4 of the 6 individuals where we were able to obtain more than 10 sequences (Fig. 2c). We conclude that cells responding to HIV-gag, CMV-pp65, and CEFT peptides contain clonal proviruses.

We next analyzed the sequences obtained from defective proviruses to determine how unique and clonal proviruses contributed to the overall enrichment observed among total HIV-1 sequences. Unique sequences were enriched in CMV-pp65 responsive cells from participants 605, 9255, 5104, and 5106 compared to total live CD4^+^ T cells in the MOG control (Fig. 3a). There was also decreased representation of unique sequences in response to some antigens in 9247 and 9241 (Fig. 3a). Otherwise, no distinct pattern emerged from the analysis of defective single viruses and there was no significant antigen dependent enrichment among unique sequences between the other individuals.

**Figure 3.**
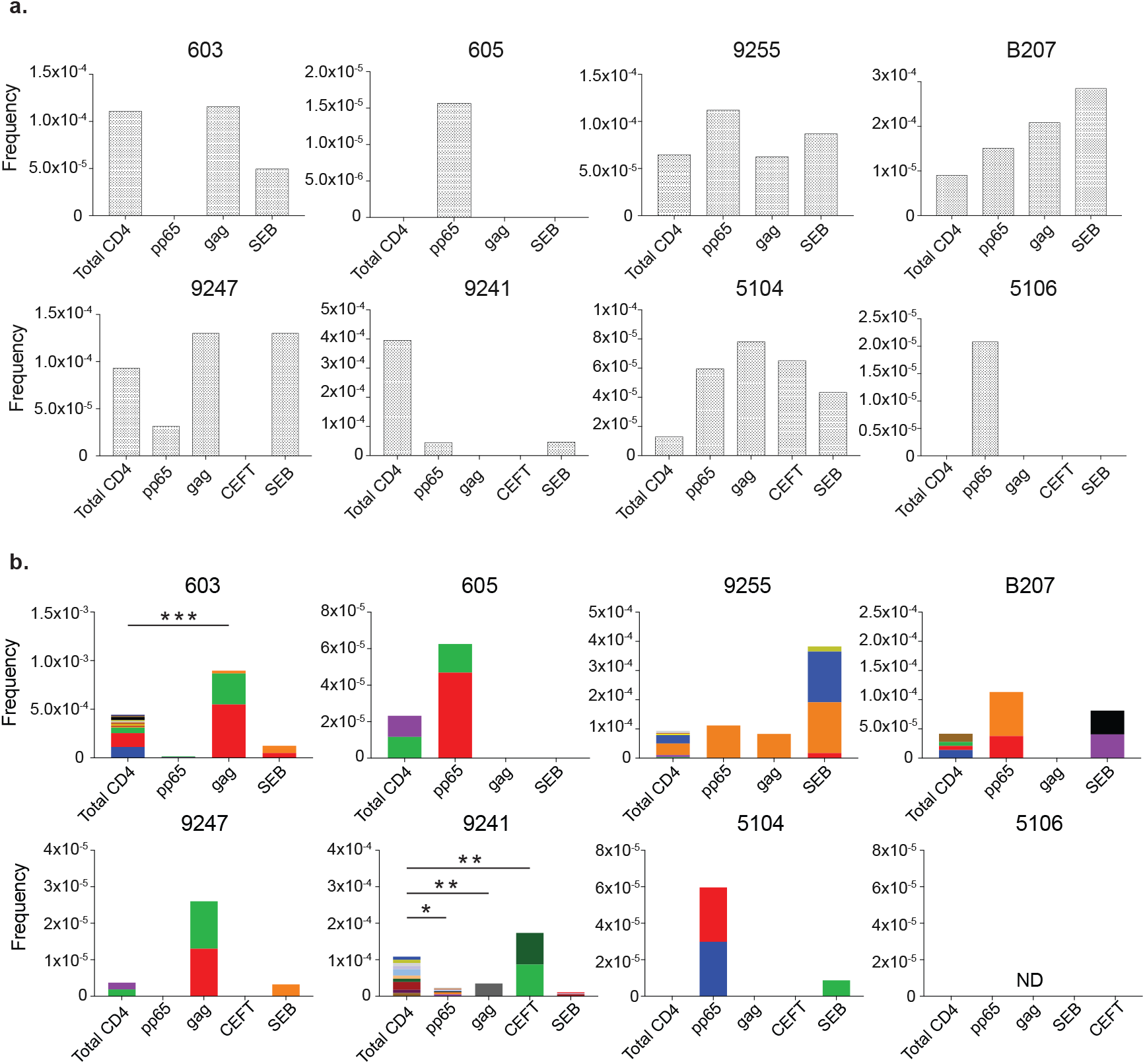
(a) Frequency of unique defective proviruses isolated from AIM+ cells, calculated by dividing number of sequences by total number of CD4^+^ T cell analyzed. (b) Frequency of clonal defective proviruses isolated from AIM+ cells, calculated by dividing number of clonal sequences by total number of CD4^+^ T cell analyzed. Each color represents a unique clone. In each donor, clones identified in more than one fraction of cells are represented by the same color. Asterisks denote a significant change in overall clonal distribution (* *P* ≤ 0.05; ** *P* ≤ 0.01; *** *P* ≤ 0.001, two-sided Fisher’s exact test).

Conversely, 7 of the 8 individuals tested showed clones of identical defective proviruses in HIV-gag, or CMV-pp65 or CEFT responsive CD4^+^ T cells (Fig. 3b). The overall enrichment of defective clonal sequences among antigen-responsive cells was often attributable to one or more specific clones. For example, in individuals 603 and 9247 the relative enrichment of defective proviruses corresponds to 2 clones isolated from HIV-gag responsive CD4^+^ T cells, whereas in 605, 5104 and B207, 2 defective clones account for the HIV-1 enrichment in CMV-pp65-responsive CD4^+^ T cells (Fig. 3b). In addition, participant 9241 showed an enrichment of 2 proviral clones in CEFT responsive cells (Fig. 3b). We conclude that expanded clones of cells harboring defective HIV-1 proviruses are responsive to peptides derived from HIV-gag, CMV-pp65 and CEFT antigens.

SEB is a superantigen that stimulates T cells expressing only a subset of T cell receptors (Llewelyn et al., 2006). Consistent with this limited specificity, we found that 4 of the 8 individuals tested (9255, B207, 9247 and 5104) had expanded or novel clones of defective proviruses in SEB-reactive CD4^+^ T cells, compared to the MOG control (Fig. 3b).

Similar analysis was also performed using intact proviruses, which as expected, were present in much smaller numbers than defective proviruses (Fig. 4, (Bruner et al., 2016; Hiener et al., 2017; Ho et al., 2013). Unique proviruses were enriched 15-fold in HIV-gag-specific CD4^+^ T cells in participant B207, while in 605, they were found in CMV-pp65 responsive cells (Fig. 4a).

**Figure 4.**
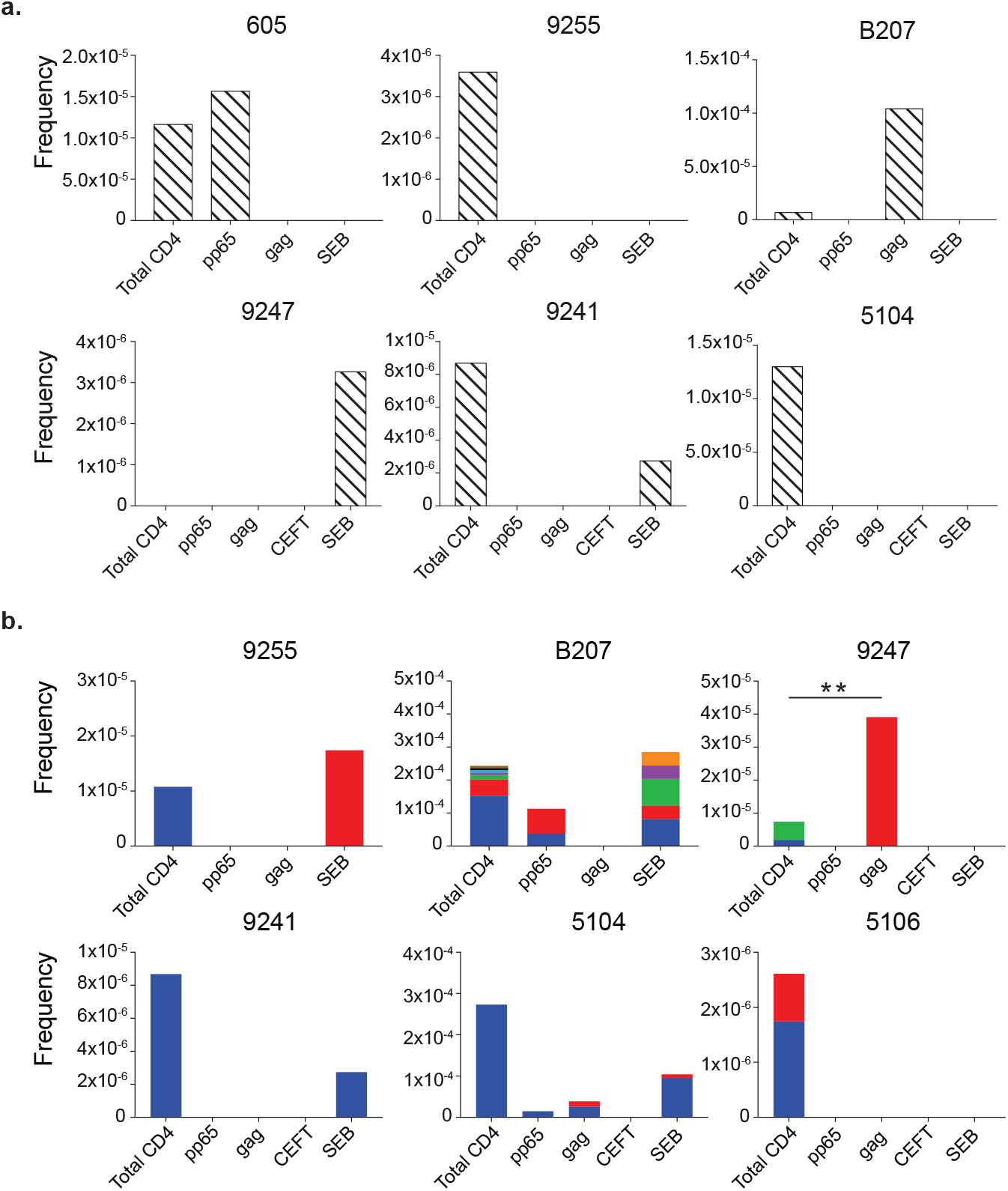
(a) Frequency of unique intact proviruses isolated from AIM+ cells. (b) Frequency of clonal intact proviruses isolated from AIM+ cells. Each color represents a unique clone. In each donor, clones identified in more than one fraction of cells are represented by the same color. Asterisks denote a significant change in overall clonal distribution (* *P* ≤ 0.05; ** *P* ≤ 0.01; *** *P* ≤ 0.001, two-sided Fisher’s exact test).

Antigen or SEB responsive CD4^+^ T cell clones harboring intact proviruses were identified in 3 and 4 of the 8 participants, respectively (Fig. 4b). In B207 and 9247 we found clones of CD4^+^ T cells responding to CMV-pp65 or HIV-gag respectively. In 9247 this activity was due to a single intact clone of identical proviruses. Although intact proviral clones were found in total CD4^+^ T cells in the MOG control in participants 9255, 9241, 5104 and 5106, they were nearly entirely absent from antigen responsive cells (Fig. 4b). However, the absence of such sequences might be due to the limited number of cells and panel of antigens assayed. We conclude that cells harboring intact proviruses can respond to peptides from HIV-1 and CMV antigens.

To determine whether the intact sequences isolated from antigen-responsive cells are replication competent, we compared the sequences obtained from antigen or SEB responsive cells to those obtained from single-cell viral outgrowth cultures (Q^2^VOA) from 5 of the individuals for whom such data was available (Cohn et al., 2018; Mendoza et al., 2018). B207 and 9247 had identical matches between Q^2^VOA sequences and proviral sequences isolated from either CMV-pp65- or HIV-gag-responsive cells (Fig. 5). Identical matches for proviruses found in SEB reactive CD4^+^ T cells were found in 9255, and B207. Finally, closely related sequences were found in the remaining individuals (Fig. 5). We conclude that clones of CD4^+^ T cells responsive to HIV-gag, CMV-pp65, and SEB harbor replication competent viruses.

**Figure 5.**
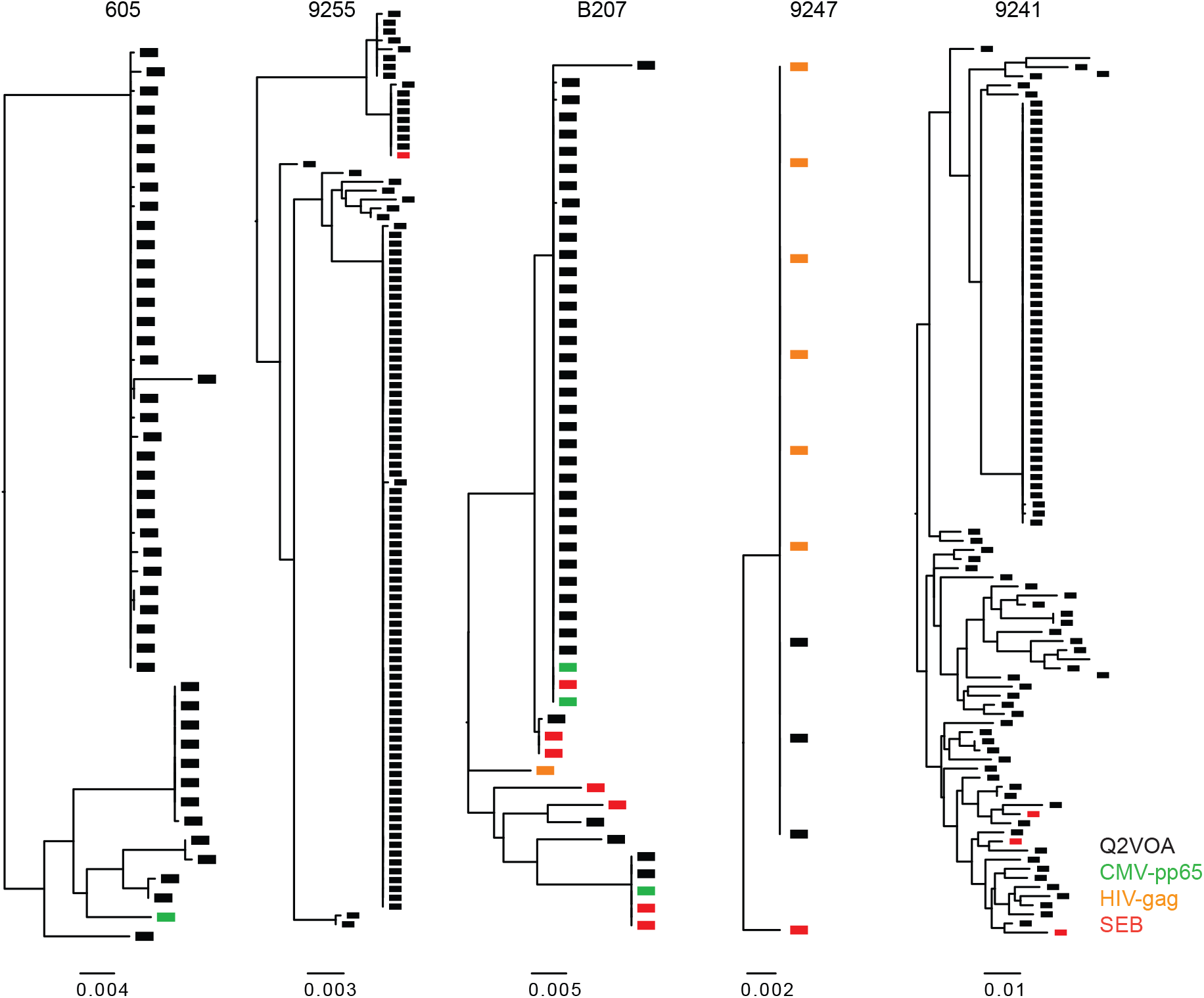
Phylogenetic tree of env sequences. Maximum likelihood phylogenetic trees of env sequences from intact viruses obtained from AIM+ cells (green: CMV-pp65, orange: HIV-gag, red: SEB) and Q^2^VOA single cell virus outgrowth cultures (black).

## Discussion

Understanding the mechanism of latent reservoir persistence is critical to developing methods for HIV-1 eradication or functional cure. We have used a limited number of antigens to test the idea that CD4^+^ T cells harboring latent proviruses can undergo clonal expansion in response to antigenic stimulation. Our results suggest a mechanism for viral persistence whereby clones of infected cells harboring latent proviruses are stimulated to divide during immune responses.

A number of ideas have been proposed to explain HIV-1 persistence. These include low-level ongoing HIV-1 replication in sanctuary sites where drug concentrations are insufficient to block virus replication (Lorenzo-Redondo et al., 2016), and infected T cell longevity (Hatano et al., 2013; Sengupta and Siliciano, 2018). Cell division was not initially considered as a mechanism of proviral persistence because the signals that induce T cell division, such as NF-κB, also tend to reactivate latent proviruses and might lead to cell death (Siliciano and Greene, 2011). However, several lines of evidence support the idea that CD4^+^ T cells harboring latent proviruses can undergo clonal expansion. Initial support for clonal expansion came from longitudinal human studies showing that ART suppressed individuals produce closely related viruses at low levels over time (Bailey et al., 2006; Tobin et al., 2005). Identical viruses were also obtained from independent cultures of latent CD4^+^ T cells in limiting dilution viral outgrowth assays suggesting that at least 50% of the circulating reservoir consists of expanded clones (Lorenzi et al., 2016; Hosmane et al., 2017; Bui et al., 2017; Simonetti et al., 2016; Wiegand et al., 2017). In addition, proviral integration site sequencing revealed collections of cells that share identical proviral integration sites (Cohn et al., 2015; Wagner et al., 2014; Maldarelli et al., 2014). Furthermore, *in vitro* studies showed that cell division and latency are not mutually exclusive (Wang et al., 2018; Hosmane et al., 2017; Pinzone et al., 2019). Finally, isolation of productively infected cells *ex vivo* (Cohn et al., 2018) and paired full-length viral sequencing and integration site analysis (Einkauf et al., 2019) provided definitive evidence for the existence of expanded clones of CD4^+^ T cells harboring intact latent proviruses.

Despite compelling evidence for CD4^+^ T cell proliferation as a mechanism for HIV-1 persistence, how latently infected cells divide without succumbing to cytopathic viral reactivation remains unknown. Possible explanations include expression of genes that favor survival by suppressing viral transcription (Cohn et al., 2018), viral integration in transcriptionally inactive regions of the genome (Craigie and Bushman, 2012), and expression of anti-apoptotic proteins such as BIRC5 (Kuo et al., 2018).

Although there is general agreement that a large fraction of the HIV-1 reservoir is comprised of clones of CD4^+^ T cells that wax and wane in circulation (Wang et al., 2018), far less is understood about what triggers clonal expansion and maintains reservoir longevity. It has been proposed that proviral integration in the vicinity of genes that control cell division, such as cancer associated genes, promotes cell growth (Maldarelli et al., 2014; Wagner et al., 2014). However, HIV-1 is preferentially integrated into highly transcribed genes (Schröder et al., 2002) which include many cancer-associated genes. Thus, it has been difficult to definitively determine whether or how integration in the vicinity of cancer related genes contributes to HIV-1 persistence (Cohn et al., 2015; Einkauf et al., 2019). Moreover, unlike transforming retroviruses that integrate into cancer genes and cause unrestricted cell growth, HIV-1 is not known to cause T cell cancers by integration.

Despite the use of a very limited number of test antigens, our work suggests that expanded clones of CD4^+^ T cells containing replication competent proviruses respond to pathogens. Their intermittent exposure to these and other antigens found in the virome and microbiome may account for the reported waxing and waning of individual clones of latently infected CD4^+^ T cells and their persistence over time. Finally, barrier tissues that are chronically exposed to antigen, such as the skin or gut, may represent reservoir sites that contribute to persistence of the reservoir by supporting ongoing expansion and contraction of CD4^+^ T cell clones harboring latent proviruses.

## Materials and Methods

### Study design and participants

All study participants were recruited by the Rockefeller University Hospital, New York, NY, and the University Hospital Cologne, Cologne, Germany, in two open-label studies (http://www.clinicaltrials.gov; NCT02825797; (Mendoza et al., 2018); NCT02588586; (Cohen et al., 2018); and NCT03571204). All participants provided written informed consent before participation in the studies. The studies were conducted in accordance with Good Clinical Practice. The protocols were approved by the US Food and Drug Administration, the Paul Ehrlich Institute (Germany), and the institutional review boards at the Rockefeller University and the University of Cologne. PBMCs were isolated from leukapheresis by Ficoll separation and frozen in aliquots. All participants were confirmed to be aviremic at the time of sample collection. The samples used in the study described here were collected before therapeutic intervention.

### CD4^+^ T cell isolation

Total CD4^+^ T cells from baseline leukapheresis were isolated from cryopreserved PBMCs by manual magnetic labeling and negative selection using the CD4^+^ T Cell Isolation Kit (Miltenyi Biotec).

### Activation-induced marker (AIM) assay

PBMCs were thawed, washed, and CD8^+^ T cells were depleted using human CD8 MicroBeads (Miltenyi, Cat# 130-045-201). CD8^+^ T cell-depleted PBMCs were cultured in 24 well plates at a concentration of 10 × 10^6^ cells/ml in RPMI 1640 supplemented with HEPES, penicillin/streptomycin and 10% human serum (Sigma). Cells were rested for 3h and then stimulated with 0.5 ug/ml of HIV-1 consensus B Gag pool, CMV peptide pool, and CEFT peptide pool (PM-HIV-CONB, PM-PP65-2, PM-CEFT-MHC-II, all from JPT Peptide Technologies) for 18h at 37°C, 5% CO_2_. A MOG peptide pool (0.5 ug/ml, PM-MOG, JPT Peptide Technologies) served as negative control. SEB-stimulated cells (0.5 ug/ml, Toxin technology, Cat# BT202) served as positive control. Cells were stained for viability with Aquavivid (Life Technologies) at 1/200 in PBS for 20 min, 4°C; incubated with human FcR Block (Miltenyi Biotec, Cat#130-059-901) at 1/70 in PBS-FBS 1% for 10 min, 4°C; and stained with antibodies to surface markers for 30 min, 4°C with Brilliant Violet Buffer (BD Biosciences, Cat# 566349) at 1/4 in PBS-FBS 1% (see Supplemental Table 3 for antibody staining panel).

### Cell sorting

AIM+ cells were gated on live, single cells CD14^−^CD19^−^CD8^−^CD4^+^ and were sorted for expression of CD69 and/or PD-L1, and/or 4-1BB. MOG stimulated cells were sorted based on CD14^−^CD19^−^CD8^−^CD4^+^ expression. Cells were sorted on a FACS Aria II flow cytometer (BD Biosciences) into tubes containing RPMI media supplemented with 10% of FBS, HEPES and Pen/Strep. MOG and SEB-treated cells served as negative and positive controls, respectively, to set gates for sorting. Cells were pelleted and cell pellets were flash frozen on dry ice and subsequently processed for DNA extraction as described below.

### Data analysis

Flow cytometric data was analyzed using FlowJo version 10.6.0 for Mac. R programming language was used to apply the Fisher’s Exact Test to evaluate whether there was a statistically significant change in the overall distribution of the clones between the Q4PCR-derived HIV-1 sequences from total CD4^+^ T cells (MOG control) and all antigen-reactive AIM+ CD4^+^ T cells.

### DNA isolation and quantification

Genomic DNA from baseline or sorted CD4^+^ T cells was isolated using phenol-chloroform (Klein et al., 2011). Briefly, CD4^+^ T cells were lysed in Proteinase K buffer (100 mM Tris, pH 8, 0.2% SDS, 200 mM NaCl, and 5 mM EDTA) and 20 mg/ml Proteinase K at 56°C for 12 h followed by genomic DNA extraction with phenol/chloroform/isoamyl alcohol (Invitrogen), non-fluorescent pellet paint (Millipore, 70748) and ethanol precipitation. DNA concentration was measured by Qubit High Sensitivity Kit (Thermo Fisher).

### Limiting dilution *gag* qPCR

To get a baseline measurement of cells containing gag+ proviruses, genomic DNA from total CD4^+^ T cells was serially diluted and assayed by gag qPCR. Based on the Poisson distribution, when <30% of qPCR reactions are positive, each positive PCR reaction has a >80% probability of containing a single copy of HIV-1 DNA (Lu et al., 2018). The selected dilution from total CD4^+^ T cells became the baseline, and genomic DNA from sorted cells was serially diluted depending on this baseline (concentrations ranged from 600 to 12.5 CD4^+^ T cells per well). DNA was assayed in a minimum of 16 reactions per concentration in a 384-well plate format using the Applied Biosystem QuantStudio 6 Flex Real-Time PCR System. HIV-1–specific primers and a probe targeting a conserved region in *gag* were used in a limiting dilution qPCR reaction (forward primer, 5’-ATGTTT TCAGCATTATCAGAAGGA-3’; internal probe, 5’-/6-FAM/CCACCCCAC/ZEN/AAGATTTAAACACCATGCTAA/3’/IABkFQ/; reverse primer, 5’-TGCTTGATGTCCCCCCACT-3’; Integrated DNA Technologies; (Palmer et al., 2003)).

Each qPCR reaction was performed in a 10 μl total reaction volume containing 5 μl of TaqMan Universal PCR Master Mix containing Rox (catalog no. 4304437; Applied Biosystems), 1 μl of diluted genomic DNA, nuclease free water, and the following primer and probe concentrations: 337.5 nM of forward and reverse primers with 93.75 nm of *gag* internal probe. *gag* qPCR conditions were 94°C for 10 min, 50 cycles of 94°C for 15 s, and 60°C for 60 s.

### NFL HIV-1 PCR (NFL1)

We used a two-step nested PCR approach to amplify NFL HIV-1 genomes. All reactions were performed in a 20 μl reaction volume using Platinum Taq High Fidelity polymerase (Thermo Fisher Scientific). The outer PCR reaction (NFL1) was performed on genomic DNA at a single-copy dilution (previously determined by *gag* limiting dilution) using outer PCR primers BLOuterF (5’-AAATCTCTAGCAGTGGCGCCCGAACAG-3’) and BLOuterR (5’-TGAGGGATCTCTAGTTACCAGAGTC-3’). Touchdown cycling conditions were 94°C for 2 min and then 94°C for 30 s, 64°C for 30 s, and 68°C for 10 min for three cycles; 94°C for 30 s, 61°C for 30 s, and 68°C for 10 min for three cycles; 94°C for 30 s, 58°C for 30 s, and 68°C for 10 min for three cycles; 94°C for 30 s, 55°C for 30 s, and 68°C for 10 min for 41 cycles; and then 68°C for 10 min (Li et al., 2006; Ho et al., 2013).

### Q4PCR

Undiluted 1-μl aliquots of the NFL1 PCR product were subjected to a Q4PCR reaction using a combination of four primer/probe sets that target conserved regions in the HIV-1 genome. Each primer/probe set consists of a forward and reverse primer pair as well as a fluorescently labeled internal hydrolysis probe as follows: PS: forward, 5’-TCTCTC GACGCAGGACTC-3’; reverse, 5’-TCTAGCCTCCGCTAGTCAAA-3’; probe, 5’/Cy5/TTTGGCGTA/TAO/CTCACCAGTCGCC/3’/IAbRQSp (Integrated DNA Technologies) (Bruner and Siliciano, 2016); *env*: forward, 5’-AGTGGTGCAGAGAGA AAAAAGAGC-3’; reverse, 5’-GTCTGGCCTGTACCGTCAGC-3’; probe, 5’/VIC/CCTTGGGTTCTTGGGA/3’/MGB (Thermo Fisher Scientific; (Bruner et al., 2016)); *gag*: forward, 5’-ATGTTTTCAGCATTATCAGAAGGA-3’; reverse, 5’-TGCTTG ATGTCCCCCCACT-3’; probe, 5’/6-FAM/CCACCCCAC/ZEN/AAGATTTAAACACCATGCTAA/3’/IABkFQ (Integrated DNA Technologies; (Palmer et al., 2003)); and *pol*: forward, 5’-GCACTTTAAATTTTCCCA TTAGTCCTA-3’; reverse, 5’-CAAATTTCTACTAATGCTTTTATTTTTTC-3’; probe, 5’/NED/AAGCCAGGAATGGATGGCC/3’/MGB (Thermo Fisher Scientific; (Schmid et al., 2010)).

Each Q4PCR reaction was performed in a 10-μl total reaction volume containing 5 μl TaqMan Universal PCR Master Mix containing Rox (4304437; Applied Biosystems), 1 μl diluted genomic DNA, nuclease-free water, and the following primer and probe concentrations: PS, 675 nM of forward and reverse primers with 187.5 nM of PS internal probe; *env*, 90 nM of forward and reverse primers with 25 nM of *env* internal probe; *gag*, 337.5 nM of forward and reverse primers with 93.75 nM of *gag* internal probe; and *pol*, 675 nM of forward and reverse primers with 187.5 nM of *pol* internal probe. qPCR conditions were 94°C for 10 min, 40 cycles of 94°C for 15 s, and 60°C for 60 s. All qPCR reactions were performed in a 384-well plate format using the Applied Biosystem QuantStudio 6 Flex Real-Time PCR System.

### qPCR data analysis

We used QuantStudio Real-Time PCR Software version 1.3 (Thermo Fisher Scientific) for data analysis. The same baseline correction (start cycle: 3, end cycle: 10) and normalized reporter signal (ΔRn) threshold (ΔRn threshold = 0.025) was set manually for all targets/probes. Fluorescent signal above the threshold was used to determine the threshold cycle. Samples with a threshold cycle value between 10 and 40 of any probe or probe combination were identified. Samples showing reactivity with two or more of the four qPCR probes were selected for further processing.

### Nested NFL HIV-1 PCR (NFL2)

The nested PCR reaction (NFL2) was performed on undiluted 1 μl aliquots of the NFL1 PCR product. Reactions were performed in a 20 μl reaction volume using Platinum Taq High Fidelity polymerase (Thermo Fisher Scientific) and PCR primers 275F (5’-ACA GGGACCTGAAAGCGAAAG-3’) and 280R (5’-CTAGTTACCAGAGTCACACAACAG ACG-3’; (Ho et al., 2013)) at a concentration of 800 nM. Touchdown cycling conditions were 94°C for 2 min and then 94°C for 30 s, 64°C for 30 s, and 68°C for 10 min for three cycles; 94°C for 30 s, 61°C for 30 s, and 68°C for 10 min for three cycles; 94°C for 30 s, 58°C for 30 s, and 68°C for 10 min for three cycles; 94°C for 30 s, 55°C for 30 s, and 68°C for 10 min for 41 cycles; and then 68°C for 10 min.

### Library preparation and sequencing

All nested PCR products from NFL2 were subjected to library preparation. The Qubit 3.0 Fluorometer and Qubit dsDNA BR Assay Kit (Thermo Fisher Scientific) was used to measure DNA concentrations. Samples were diluted to a concentration of 10–20 ng/μl. Tagmentation reactions were performed using 1 μl of diluted DNA, 0.25 μl Nextera TDE1 Tagment DNA enzyme (catalog no. 15027865), and 1.25 μl TD Tagment DNA buffer (catalog no. 15027866; Illumina). Tagmented DNA was ligated to unique i5/i7 barcoded primer combinations using the Illumina Nextera XT Index Kit v2 and KAPA HiFi HotStart ReadyMix (2X; KAPA Biosystems) and then purified using AmPure Beads XP (Agencourt). 384 purified samples were pooled into one library and then subjected to paired-end sequencing using Illumina MiSeq Nano 300 V2 cycle kits (Illumina) at a concentration of 12 pM.

### HIV-1 sequence assembly and annotation

HIV-1 sequence assembly was performed by our in-house pipeline (Defective and Intact HIV genome Assembler), which is capable of reconstructing thousands of HIV genomes within hours via the assembly of raw sequencing reads into annotated HIV genomes. The steps executed by the pipeline are described briefly as follows. First, we removed PCR amplification and performed error correction using clumpify.sh from BBtools package v38.72 (http://sourceforge.net/projects/bbmap/). After, a quality-control check was carried out by Trim Galore package v0.6.4 to trim Illumina adapters and low-quality bases. (https://github.com/FelixKrueger/TrimGalore). We also used bbduk.sh from BBtools package to remove possible contaminant reads using HIV genome sequences, obtained from Los Alamos HIV database, as a positive control. We used a k-mer based assembler, SPAdes v3.13.1, to reconstruct the HIV-1 sequences. The longest assembled contig was aligned via BLAST to a database of HIV genome sequences, obtained from Los Alamos, in order to set the correct orientation. Finally, the HIV genome sequence was annotated by aligning against Hxb2 using BLAST. Sequences with double-peaks, i.e., regions indicating the presence of two or more viruses in the sample (cut-off consensus identity for any residue < 70%), or samples with a limited number of reads (empty wells ≤500 sequencing reads) were omitted from downstream analyses. In the end, sequences were classified as intact or defective. Defective sequences were subdivided into more specific classifications according to their sequence structure: Major Splice Donor (MSD) Mutation, Non-functional, or Missing Internal Genes.

### Clone analysis for intact and defective sequences

Clones were defined by aligning sequences of each classification (Intact, MSD Mutation, Non-functional, and Missing Internal Genes) to HXB2 and calculating their Hamming distance. Sequences having a maximum of three differences between the first nucleotide of Gag and the last nucleotide of Nef (reference: HXB2) were considered members of the same clone.

### Data availability

Data is available upon request.

## Acknowledgements

We thank all study participants who devoted time to our research; The Rockefeller University Hospital and nursing staff, members of the Nussenzweig laboratory, Edward R. Kastenhuber for computational analysis, Don Ganem and Peter Kim for helpful discussions, the Biohub lab management team, Maša Jankovic for laboratory management, and especially to Zoran Jankovic for his unwavering support and for running the lab with enormous dedication and commitment. Research reported in this publication was supported by the Chan Zuckerberg Biohub, the National Institute Of Allergy And Infectious Diseases of the National Institutes of Health (grant UM1AI126611 supported L.B. Cohn, and grants 1UM1 AI100663 and R01AI129795; to M.C. Nussenzweig); the Bill and Melinda Gates Foundation (Collaboration for AIDS Vaccine discovery grants OPP1092074, OPP1124068, and OPP1168933) and the Einstein–Rockefeller–CUNY Center for AIDS Research (grant 1P30AI124414-01A1); and the Robertson Fund. C. Gaebler was supported by the Robert S. Wennett Post-Doctoral Fellowship, in part by the National Center for Advancing Translational Sciences (National Institutes of Health Clinical and Translational Science Award program, grant UL1 TR001866), and by the Shapiro-Silverberg Fund for the Advancement of Translational Research. The content is solely the responsibility of the authors and does not necessarily represent the official views of the National Institutes of Health. M.C. Nussenzweig is a Howard Hughes Medical Institute Investigator.

## Author Contributions

P. Mendoza, M.C. Nussenzweig and L.B. Cohn designed the research. P. Mendoza, J.R. Jackson, C. Gaebler, and L.B. Cohn performed the research. M.C. performed study subject recruitment and oversaw sample collection. P. Mendoza, T.Y. Oliveira, V. Ramos, M. Jankovic, M.C. Nussenzweig and L.B. Cohn analyzed the data. P. Mendoza, M.C. Nussenzweig, and L.B. Cohn wrote the manuscript.

---

**Supplemental Figure 1.**
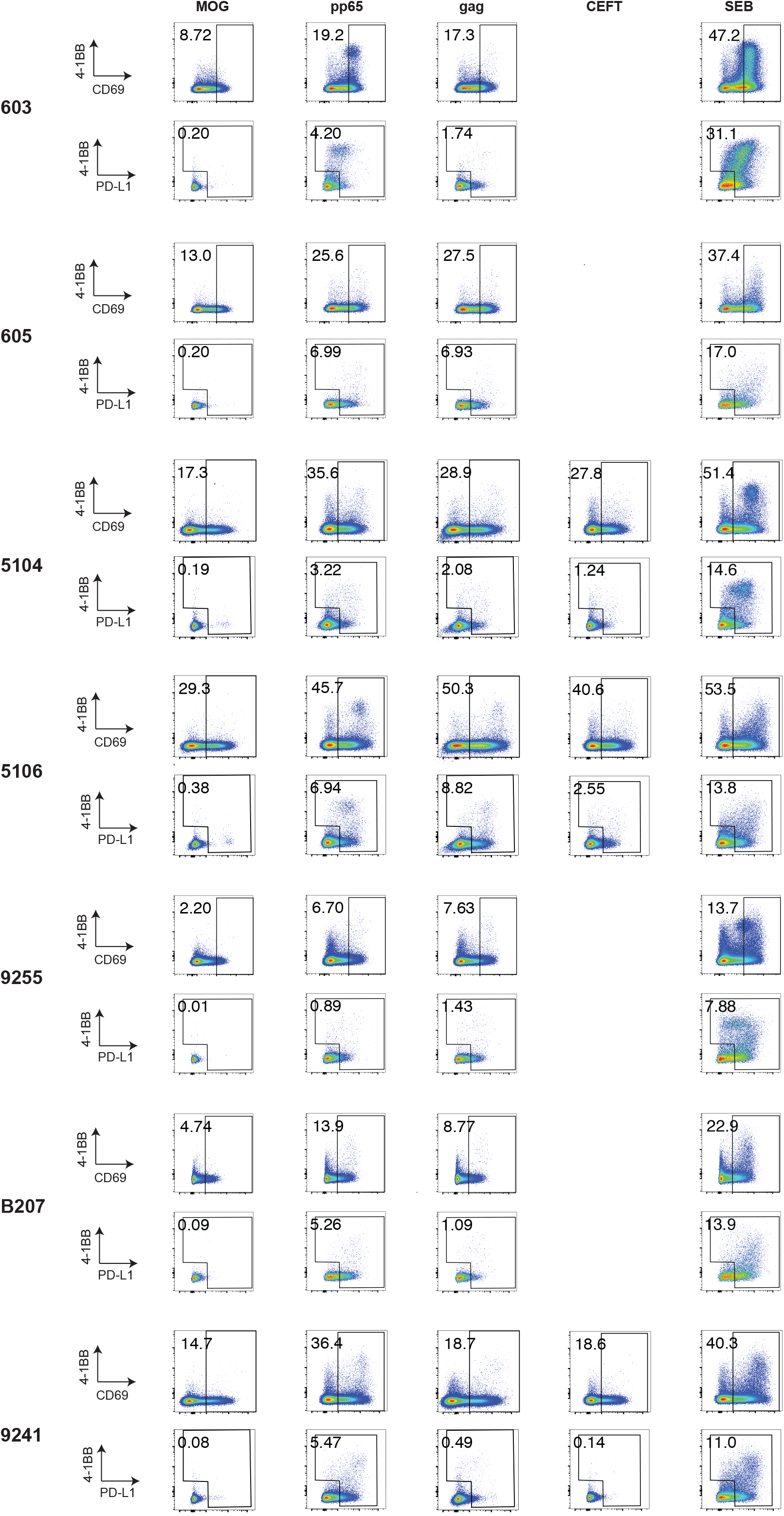
CD4^+^ T cells were stimulated for 18h with either peptide pools from myelin oligodendrocyte glycoprotein (MOG; control), CMV-pp65, HIV-gag, or CMV, EBV, influenza and tetanus peptides (CEFT) before staining. CD4^+^ T cells gated on CD69-positive cells followed by gating on PD-L1 positive and/or 4-1BB positive cells. Each AIM assay staining was performed twice on each donor. Numbers represent the percentage of total CD4^+^ T cells within each gate.

**Supplemental Table 1.** Individual participant demographics and baseline clinical characteristics

| ID | Age | Gender | Race | Year HIV-1 Dx | Year ART initiation | Uninterr. ART (yrs) | Viral load (copies/ml) | CD4 T cell count | Reported nadir | ART regimen* |
| --- | --- | --- | --- | --- | --- | --- | --- | --- | --- | --- |
| **9241** | 40 | M | White/Hisp | 2012 | 2013 | 5 | < 20 | 515 | 500 | EVG/cobi/TDF/FTC |
| **9247** | 31 | M | Black | 2012 | 2012 | 6 | < 20 | 728 | 600 | EVG/cobi/TAF/FTC |
| **9255** | 30 | M | White | 2013 | 2014 | 4 | < 20 | 1360 | 779 | EVG/cobi/TAF/FTC |
| **B207** | 48 | M | White/Hisp | 2006 | 2007 | 11 | < 20 | 724 | 50 | EVG/TDF/FTC |
| **603** | 45 | M | White/Hisp | 2003 | 2005 | 12 | < 20 | 693 | 300 | EFV/TDF/FTC |
| **605** | 38 | M | White/Hisp | 2001 | 2002 | 15 | 20 | 524 | 372 | RPV/TDF/FTC |
| **5104** | 35 | M | Black | 2011 | 2011 | 7 | < 20 | 606 | 400 | BIC/TAF/FTC |
| **5106** | 31 | M | Black | 2012 | 2012 | 6 | < 20 | 671 | > 350 | EVG/cobi/FTC/TDF |

\* EVG - elvitegravir, cobi - cobicistat, TDF - tenofovir disoproxil fumarate, FTC - emtricitabine, RPV - rilpivirine, EFV - efavirenz, TAF - tenofovir alafenamide fumarate, BIC - bictegravir.

**Supplemental Table 2.**
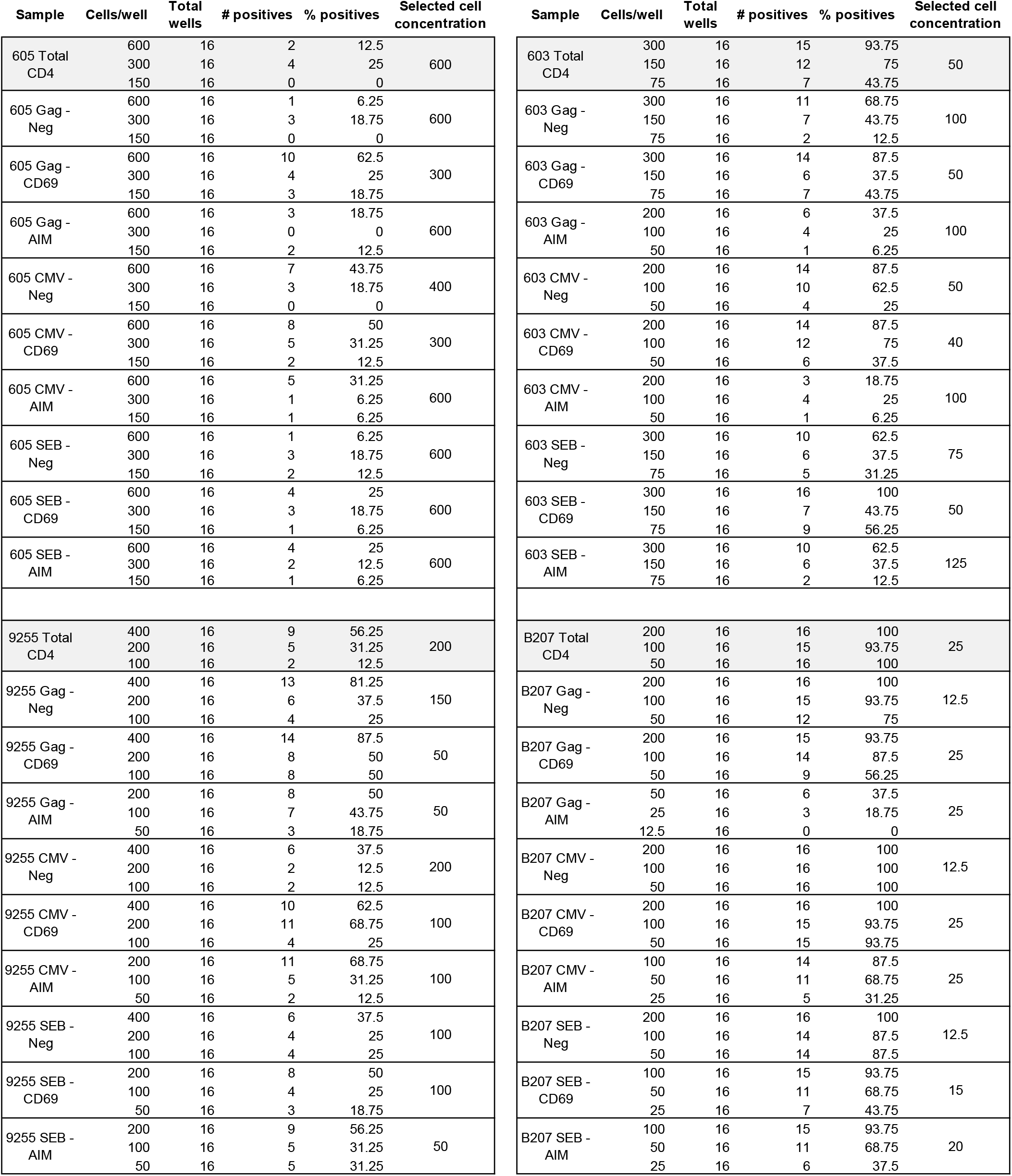
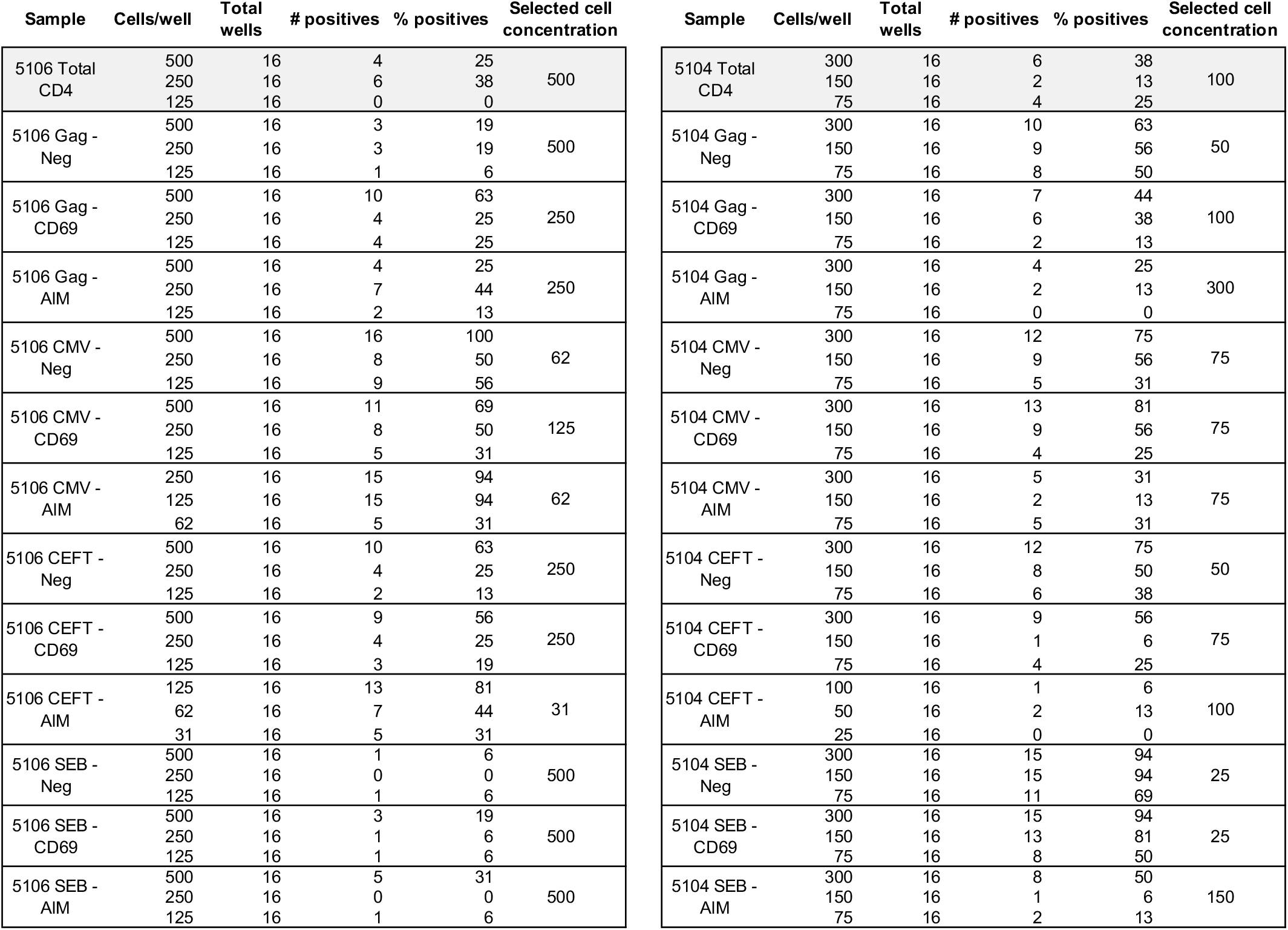
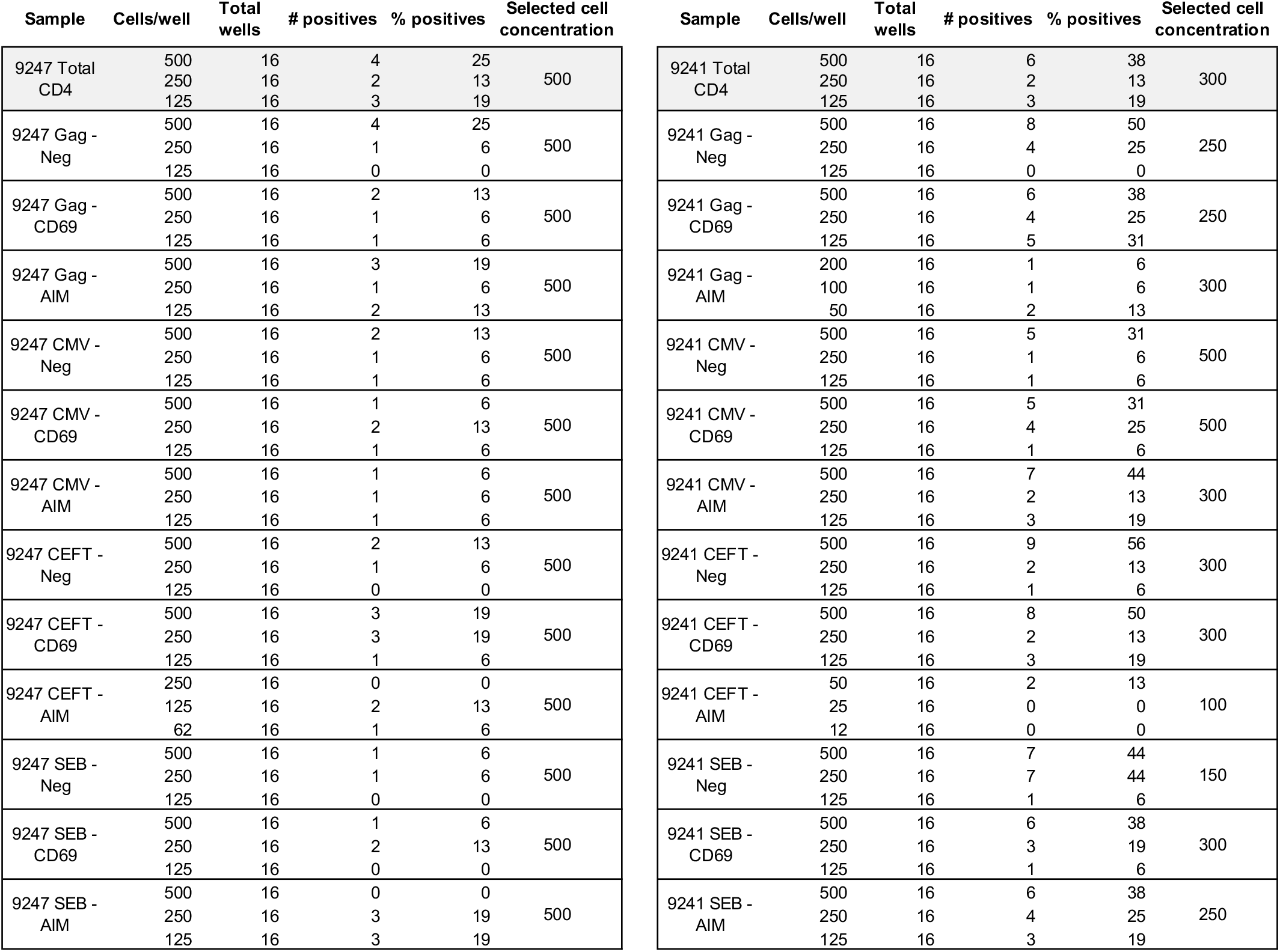
HIV-1 gag DNA enrichment in the different sorted populations. HIV-1 gag DNA was detected by qPCR with a gag probe testing 3 cellular DNA dilutions, based on previous Q4PCR analysis of these participants. A total of 16 wells for each dilution was analyzed and the number of cells per well was determined based on cellular DNA concentration. Final cell concentration for Q4PCR in each sample was determined by the dilution with less than 30% HIV-1 gag positives wells to achieve a single virus dilution.

**Supplemental Table 3.** List of antibodies used in flow cytometry for cell sorting and identification of antigen-reactive cells.

| **Marker** | **Fluorochrome** | **Clone** | **Supplier** | **Cat. No.** | **Dilution** |
| --- | --- | --- | --- | --- | --- |
| CD8 | BV711 | RPA-T8 | Biolegend | 301044 | 1/50 |
| PD-L1 | BV421 | 29E2A3 | Biolegend | 329714 | 1/20 |
| 4-1BB | PE-Cy7 | 4B4-1 | Biolegend | 309818 | 1/20 |
| CD14 | BV510 | M5E2 | Biolegend | 301842 | 1/33 |
| CD19 | BV510 | H1B19 | Biolegend | 302242 | 1/33 |
| CD4 | APC | RPA-T4 | Biolegend | 300514 | 1/133 |
| CD69 | BV650 | FN50 | Biolegend | 310934 | 1/28 |

